# Adaptive recruitment of cortex-wide recurrence for visual object recognition

**DOI:** 10.1101/2025.10.17.682937

**Authors:** Pablo Oyarzo, Johannes J.D. Singer, Kohitij Kar, Diego Vidaurre, Radoslaw M. Cichy

## Abstract

Theories of the neural mechanism underpinning rapid recognition debate whether it relies solely on a feedforward sweep through the ventral stream or instead requires recurrent processing, possibly engaging additional brain regions. Here we directly tested the “adaptive recurrence hypothesis”, that attempts to unify these disparate views by proposing that additional recurrent cortical resources beyond the visual stream are recruited when feedforward processing alone is insufficient to solve object recognition. To investigate this hypothesis, we contrasted functional MRI (fMRI) and electroencephalography (EEG) responses to compare neural responses to images that are equally well recognized by humans, but that differ in whether they could be solved by a feedforward deep neural network; a computational proxy for ventral stream feedforward processing. We found that when feedforward processing in the ventral visual stream is insufficient, additional parieto-frontal networks are rapidly and transiently recruited, representationally reconfiguring the ventral visual stream. Our results reveal that object recognition flexibly adapts through fast, cortex-wide recurrence, providing a unifying framework for competing theories of visual recognition.

## Introduction

Humans recognize objects rapidly and effortlessly even under challenging viewing conditions ^1,2^. A key neural substrate supporting this ability is the ventral visual stream ^3,4^, a hierarchically organized cortical pathway ending in inferior temporal cortex (IT), where object representations that are independent of viewing conditions emerge ^5–8^. Because object recognition unfolds rapidly ^9^, and IT neurons contain linearly decodable object identity information already within 100-150ms ^10^, prominent theories of core object recognition propose that a single feedforward sweep along the ventral visual stream is sufficient to form a robust object representation ^1^. In contrast, alternative theories propose that recurrent activity (both within and beyond the ventral stream) plays a critical role in object recognition. This notion is consistent with strong recurrent anatomical connections both within the ventral visual stream, and to all cortical lobes^11–13^. Furthermore, multiple recent findings demonstrate that recurrent activity, both within ^14–16^ and beyond ^17–20^ the ventral visual stream, contributes to object recognition by encoding object information and modulating ventral visual stream activity.

The thesis of this paper is that these contrasting theories can be reconciled by the idea that additional object recognition resources are adaptively recruited when object recognition cannot be achieved through feedforward processing within the ventral visual stream alone. The adaptive recurrence hypothesis posits that when object recognition is easy, feedforward processing within the ventral visual stream suffices to generate robust object representations. In contrast, when recognition is challenging, recurrent computations requiring additional processing time, and additional resources beyond the ventral stream are required. This graded account is supported by two kinds of evidence. First, object representations emerge slower in cluttered scenes than in uncluttered scenes ^21,22^, and for degraded compared to intact images ^16,23^. Second, ventromedial prefrontal cortex was shown to impact the ventral visual stream specifically when viewing conditions are challenging ^17^.

Directly testing the adaptive recurrence hypothesis requires contrasting brain activity for images that the ventral visual stream can resolve in a feedforward pass versus those it cannot. Here, we leverage the methodological innovation from Kar et al. ^24^, who compared feedforward convolutional neural networks (ffCNN) with primate behavior to identify images that are behaviorally matched but differ in feedforward solvability. This approach treats ffCNNs as computational proxies for feedforward processing in ventral visual stream, based on the observation that ffCNNs better predict ventral visual stream activity for clean, canonical images but deteriorate in its alignment under challenging viewing conditions that are thought to rely on additional and recurrent processes in the brain. Object images that humans recognize accurately, but a ffCNN fails to classify, are deemed challenging, therefore needing computations that the ffCNN lacks when compared to humans. In contrast, images for which ffCNN and humans perform with comparable accuracy, which we use as a control condition, are considered solvable by feedforward processing in the ventral stream alone.

To resolve the underlying neural dynamics in space and time, we recorded human neural activity using functional MRI (fMRI; for spatial specificity) and electroencephalography (EEG; for temporal specificity) while participants viewed selected challenge and control images. We show that while control images primarily engage the ventral visual stream, challenge images elicit broader cortical engagement of a network including frontal and parietal regions, accompanied by cortex-wide representational reconfiguration. Further, challenge images are solved slower than control images, indexing additional recurrent neural dynamics at interim processing stages absent for control images. Together our results support the adaptive recurrence hypothesis: additional cortical resources for object recognition must be adaptively engaged when feedforward processing within the ventral stream is insufficient, and characterize the associated neural dynamics in time, space, and network configuration.

## Results

To construct a stimulus set that selectively engages object recognition mechanisms beyond feed-forward processing in the ventral visual stream (as approximated by a ffCNN), we followed the rationale of Kar et al. ^24^. We started with a large candidate stimulus set of 1320 unique images, generated by rendering ten different objects (**Fig. 1a left**) in 132 distinct configurations (e.g. variations in rotation and size) onto naturalistic backgrounds, and naturalistic images ^25^. For each image, we obtained categorization performance (*d*′) from both humans ^24^ and a ffCNN (AlexNet; ^26^) (**Fig. 1a right**). Based on the difference between human and DNN performance, we identified images whose performance was either similar or divergent (**Fig. 1b**). Images with similar human and DNN performance (|Δ*d*′| < 0.4) were labeled control (color-coded blue). Control images are defined as those that are assumed to be reliably recognized through feedforward processing within the ventral visual stream alone. In contrast, images for which humans outperformed the DNN by more than 1.5 *d*′ units were labeled *challenge* (color-coded red). Challenge images are therefore defined as those that are assumed not be recognized through feedforward processing within the ventral visual stream alone, needing additional cortical resources. We created two final sets of 121 images each by pairing each challenge image with a control image closely matched in human performance (**Fig. 1c,d**).

**Figure 1.**
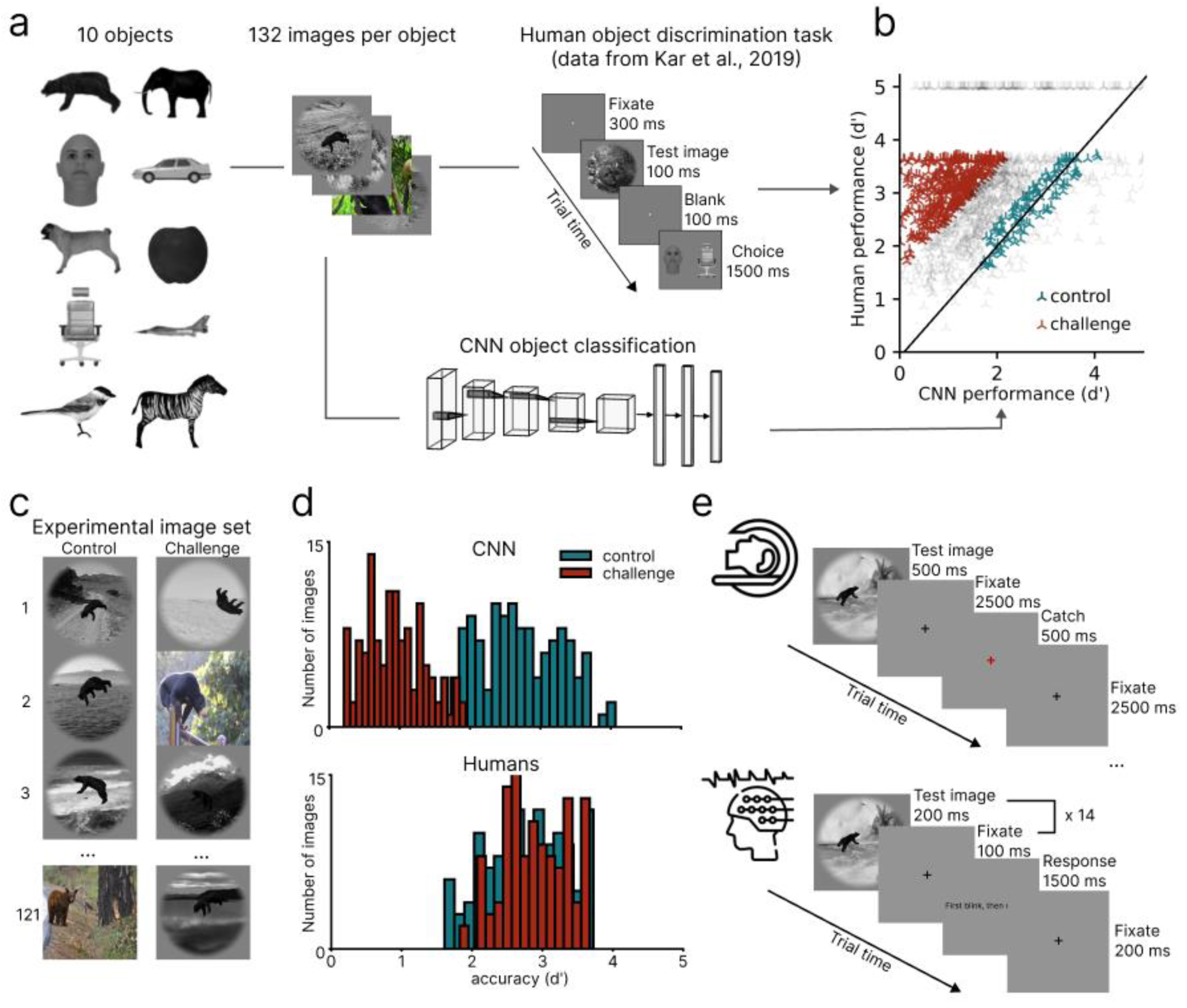
Experimental design. **a.** Selection of control and challenge images. We used a dataset of images created by rendering 10 different objects in different views (i.e. size, pose) on naturalistic backgrounds (132 images per object). The image data set was accompanied by object recognition performance (d’) estimates for humans and a feedforward convolutional neural network (CNN; AlexNet ^26^). **b.** Scatter plot comparing human and CNN performance. Each point represents a single image. Control images are defined as those with a low human–CNN performance difference (|Δd′| < 0.4; blue), while challenge images are defined as those with a high difference favoring humans (Δd′ > 1.5; red). **c.** Representative examples of control and challenge images from the final experimental set, created by selecting 121 behaviorally matched image pairs from the larger pool in (b). **d.** Performance score distributions for CNN (top) and human (bottom) across the final image set. Control (blue) and challenge (red) images were selected to minimize human performance differences (via Kuhn-Munkres algorithm), resulting in a significant difference for CNNs (u(120) = 41, p < 0.001) but not for humans (u(120) = 8155, p = 0.13). **e.** Trial structure of the fMRI (top) and EEG (bottom) experiments.

We collected fMRI (N=31) and EEG (N=34) responses to both image sets (**Fig. 1e**) from different participant groups. To minimize task-related confounds, participants performed an orthogonal task unrelated to image content during the presentation of the images. This design ensures that any differences identified between challenge and control images reflect automatic visual processing rather than interactions with explicit task demands. We then performed multivariate pattern analysis to characterize neural object representations, leveraging the fMRI to map their spatial distribution and the EEG to track their temporal dynamics.

### Challenge images engage cortex-wide resources beyond the ventral visual stream

We first characterized the spatial distribution of object representations underlying the processing of challenge and control images using the fMRI data. We predicted that while both challenge and control images are processed in the ventral visual stream ^1,2^, challenge images are processed in additional brain regions ^17–19^, indicating the engagement of additional resources. We assessed an a priori defined set of 11 regions of interest (ROIs) ^27^, comprising five core nodes of the ventral visual stream (i.e., EVC, POC, V4, LOC, IT; for definition of abbreviations see figure caption). We also considered regions anatomically connected to (and functionally interacting with) the ventral visual stream ^28,29^. These included one temporal (i.e., mTC) and five frontal ROIs (i.e., PMC, mPFC, OFC, vlPFC and dlPFC). We began by decoding object identity from multivariate activity patterns evoked by control and challenge images, using decoding accuracy as an index of explicit object representations (**Fig 2a, left**) ^30,31^. We assessed significance using non-parametric sign-flip tests (N = 31, 100,000 permutations, *p* < 0.05, Bonferroni-corrected by number of ROIs (11)).

**Figure 2.**
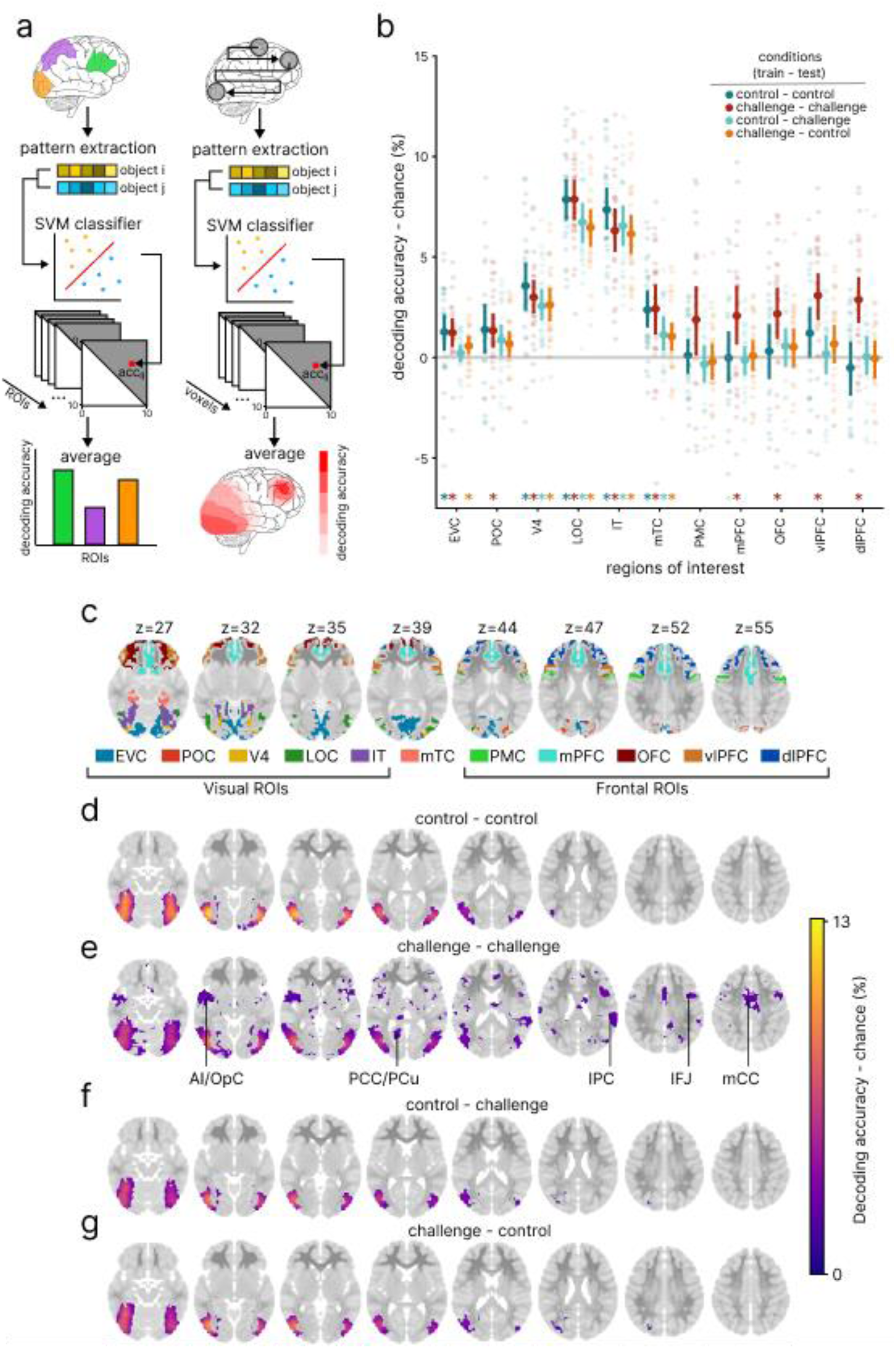
Spatial distribution of object information. **a.** Schematic of the object decoding procedure applied to fMRI. Left: ROI-based classification using activation patterns from predefined anatomical regions. Right: spatially unbiased searchlight decoding. Both analyses were performed separately for control and challenge images using a cross-validated SVM classifier. **b.** Mean object decoding accuracies (± SEM) per ROI. We targeted 11 regions in total: early visual cortex (EVC), parieto-occipital cortex (POC), V4, lateral occipital complex (LOC), inferior temporal cortex (IT), medial temporal cortex (mTC), pre-motor cortex (PMC), medial prefrontal cortex (mPFC), orbitofrontal cortex (OFC), ventrolateral prefrontal cortex (vlPFC), and dorsolateral prefrontal cortex (dlPFC). Conditions are color-coded by training–testing assignment of data to the SVM classifier. Asterisks indicate significance (p < 0.05, Bonferroni-corrected; one-sided sign-flip test across participants). **c.** Anatomical slices in MNI space used for visualizing searchlight results (see d), overlaid with color-coded ROIs used in the ROI-based decoding analysis (see b). **d–g.** Searchlight decoding results showing voxels with significant object decoding across training→testing conditions: d) control→control, e) challenge→challenge, f) control→challenge, g) challenge→control. Color scale reflects decoding accuracy; maps are thresholded at p < 0.05 using the Benjamini–Yekutieli FDR correction.

The results revealed two key insights (**Fig. 2b**). First, object information increased for both control (indicated in blue) and challenge images (red) across ventral visual cortex, peaking in high-level areas (i.e., V4, IT, LOC), and extending into the mTC. Decoding accuracy did not significantly differ between challenge and control images (paired tests; *p* > .05). This pattern is consistent with the established role of high-level ventral regions in representing object information ^1,32^, and suggests a role of mTC in object processing ^28,33^. Second, in frontal regions (mPFC, OFC, vlPFC, dlPFC), object information was decodable for challenge images (one-sample permutation test, *p* <. 001), but not for control images (*p* >. 05). This suggests that frontal cortex is additionally – and automatically – recruited alongside the ventral stream during challenge image processing.

Given that high-level ventral visual regions contained comparable object information for both image types (consistent with their role as the readout region for human visual categorization behavior ^34,35^), and that behavioral performance was equated across stimulus sets by design, we hypothesized that ventral object representations are similarly structured across conditions. Furthermore, given the exclusivity of object information in frontal cortex for challenge images, we hypothesized that frontal object representations are specific to the challenge condition.

To test these predictions, we conducted cross-decoding: classifiers were trained to predict object category from responses to control images and tested on responses to challenge images (Fig. 2b, cyan) and vice versa (orange). The results, shown in **Fig. 2b**, confirmed both predictions. In all regions where object information was present for both challenge and control images (i.e. V4, IT, LOC), cross-decoding was significant, indicating similar representations. In contrast, in frontal region there was no significant effect in cross-decoding.

To determine the presence of object representations beyond the a priori defined ROIs in a spatially unbiased fashion, we used searchlight decoding (**Fig. 2a**, right: ^36,37^) in normalized MNI space (**Fig 2c**, ^38^). We assessed significance using (voxel-wise Wilcoxon signed-rank tests, Benjamini–Yekutieli ^39^ corrected for FDR across voxels, *q* < 0.05). For control images (**Fig. 2d**) this confirmed the ROI-based results, with significant decoding accuracy largely restricted to ventral visual regions. For challenge images (**Fig. 2e**) we observed significant decoding in regions overlapping with predefined ROIs, but also in an extended brain-wide network. This included occipito-parietal regions such as the posterior cingulate and precuneus (PCC/PCu) and the inferior parietal cortex (iPC), fronto-temporal regions including the anterior insula and opercular cortex (AI/OpC) and the inferior frontal junction (IFJ), and the mid-cingulate cortex (mCC) in the midline region. Cross-decoding (**Fig. 2f,g**) revealed effects only in high-level ventral visual stream thus confirming the ROI-based results.

In sum, our results reveal a differential spatial distribution of object representations across challenge and control images. For control images, object representations were confined to the ventral visual stream. In contrast, challenge images engaged an additional, extended cortical network encompassing frontal, temporal, and parietal regions. This pattern indicates that the automatic recruitment of additional cortical resources when feedforward ventral stream processing is insufficient to support object recognition.

### Cortical networks exhibit reorganized representational structure for challenge images

The observation of an automatic recruitment of additional cortical resources beyond the ventral visual stream for challenge images is consistent with two basic yet fundamentally different views. One is that this recruitment is best described as an additive process, on top of a fixed and predetermined state of the ventral visual stream. Another is that of a reconfiguration of object representations at the cortical network level, including changes within the ventral stream itself. To arbitrate between these two mechanisms, we applied representational similarity analysis (RSA ^40^) across the set of 11 ROIs to compare the network structure of representational geometries when processing challenge versus control images (**Fig. 3a**). For each of the 11 ROIs, we computed representational dissimilarity matrices (RDMs) separately for challenge and control images, capturing each region’s representational geometry. Each RDM quantifies pairwise dissimilarities in multivariate activation patterns across all object pairs. We then compared the RDMs across all ROIs pairs, resulting in two 11×11 network similarity matrices (NSMs) —one for control (**Fig. 3b**) and one for challenge (**Fig. 3c**) images—, where each cell reflects the similarity between the representational geometries of two regions. We assessed significance of region similarities in NSMs using sign-flip permutation tests (N = 31, 100,000 permutations, *p* < 0.05, Bonferroni-corrected across ROI pairs (55).

**Figure 3.**
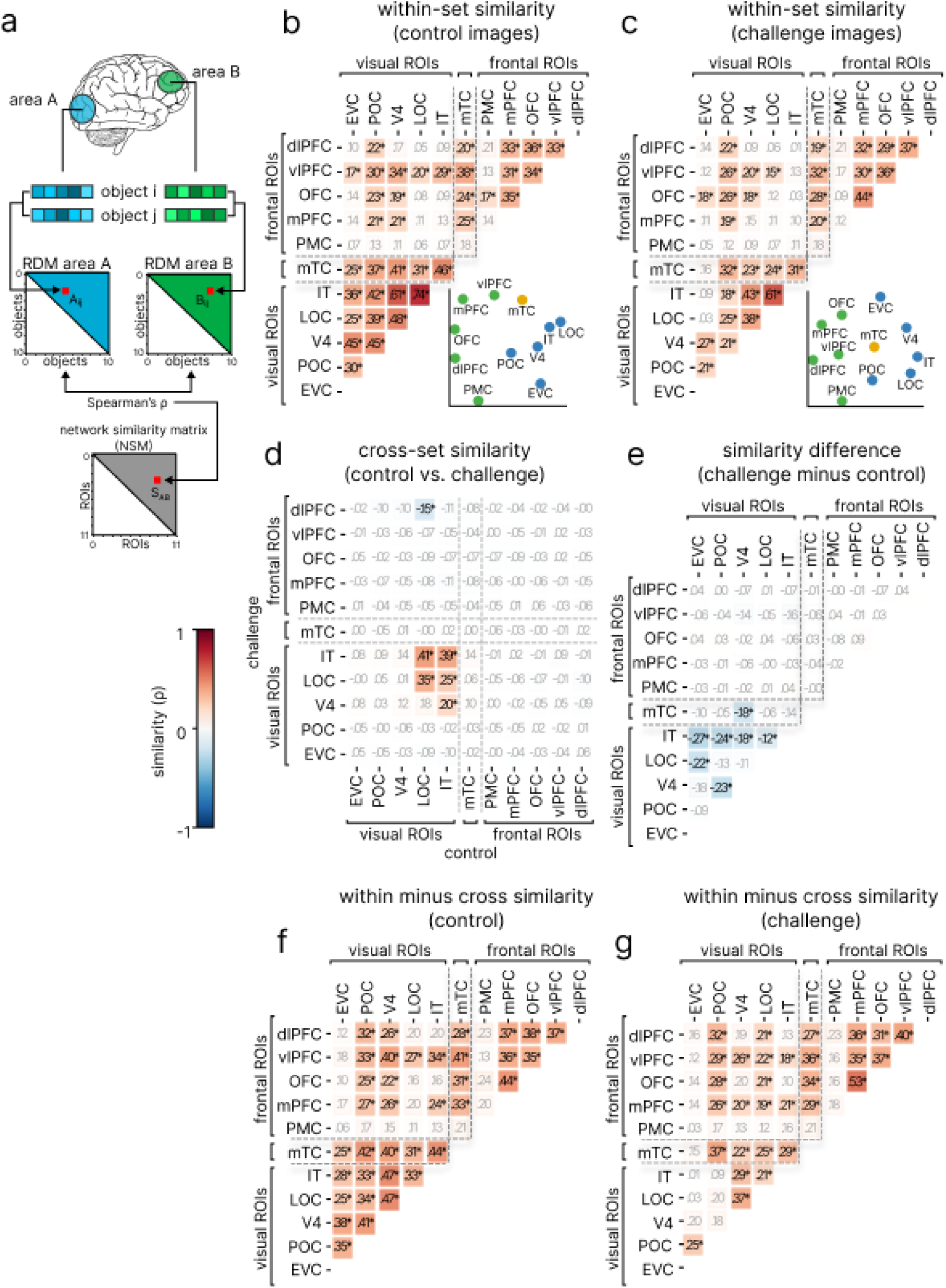
Representational reconfiguration of cortical networks. **a.** Schematic of the analysis pipeline. For each ROI and image set, we formed representational dissimilarity matrices (RDMs) by aggregating decision-value-weighted decoding accuracies across all object pairs (e.g. i,j). We quantified similarity between region-specific RDMs (e.g. A and B) using Spearman correlation, assigning the similarity values (S_AB_) to a network similarity matrix (NSM) indexed in rows and columns by ROIs compared. **b.** NSM for control images. Inset: 2D multidimensional scaling (MDS) of the NSM. ROIs are color-coded by macro-anatomical group: blue = visual regions, yellow = middle temporal cortex (mTC), green = frontal regions. **c.** NSM for challenge images. Inset as in (b) **d.** NSM computed across control and challenge image sets. **e.** Difference between NSMs for challenge (c) minus control (b) images. **f.** control NSM (b) minus cross-set (e) NSM. **g.** challenge NSM (c) minus cross-set (e) NSM. **For all panels:** Color scales represent Spearman correlation coefficients. Significant values are shown in bold and marked with an asterisk (p < 0.05, Bonferroni-corrected).

A first qualitative inspection revealed a common, basic pattern in both the challenge and control NSMs: frontal and visual regions (including mTC) clustered into two distinct groups, consistent with their respective anatomical proximity and functional organization (for visualization by multidimensional scaling see insets in **Fig. 3b,c**). To directly and quantitatively determine commonalities in representational network relations when processing challenge and control images, we computed a cross-condition NSM (**Fig. 3d**). Significant similarities were largely limited to high-level ventral regions IT and LOC (*p*s < .01), consistent with the ROI-based cross-decoding analysis (**Fig. 2b**). This indicates that high-level ventral visual cortex, specifically, maintains similar representational geometries for challenge and control images. Surprisingly, we observed significant dissimilarity between dlPFC representations for challenge images and LOC representations for control images (*p* < .01). This negative relationship might indicate an antagonistic or corrective role for dlPFC over LOC in resolving mismatched object information, potentially facilitating disambiguation of unclear configurations ^41,42^ or refining categorical boundaries ^19,20^. However, further inspection of the challenge and control NSMs revealed differences between challenge and control NSMs. Notably, EVC showed significant similarity with IT and LOC for control images (**Fig. 3b**, *p*s < .01), but not for challenge images (**Fig. 3c**, *p* = .99). More broadly, similarity among visual regions was reduced for challenge (**Fig. 3c**) compared to control (**Fig. 3b**) images. This pattern was confirmed by the subtraction of the control from the challenge NSM (**Fig. 3e**), revealing significant reductions in similarity between IT and lower visual regions (EVC, POC, V4 and LOC), and between V4 and both POC and mTC (*p* < .05). These results indicate a reconfiguration of representational geometries in visual cortex, reducing the representational similarity across visual regions for challenge images. Finally, we subtracted the cross-condition NSM from the within condition NSMs (**Fig. 3f,g**), directly quantifying condition specific differences versus their similarities in NSM structure. We observed effects for all regions, including high-level ventral visual cortex, with the exception of PMC. This indicates that processing challenge versus control images engages cortex-wide networks in distinct ways, supporting the notion of cortex-wide reconfiguration of representational relations.

Together, these results reveal that the recruitment of additional resources by challenge images is mediated by a representational reconfiguration across the brain, with high-level ventral visual cortex maintaining partially stable object representations.

### Object representations emerge later for challenge than for control images

We determined the temporal dynamics with which object representations emerge for challenge and control images. Based on our fMRI results and previous work ^24^, we predicted that object information would emerge later for challenge than for control images, reflecting the need for additional computations to yield correct object representations. To test this, we performed time-resolved decoding of object identity from EEG electrode activity patterns spanning -100 to +500 ms relative to stimulus onset. We report statistical significance assessed using sign-flip permutation tests across subjects (100,000 permutations, *p* < .05, Bonferroni corrected across time points).

The decoding time courses are shown in **Fig. 4a** for challenge (blue) and control images (red), along with their difference (yellow); peak magnitudes and latencies, along with their 95% confidence intervals, are reported in **Supplementary Table 1** and **Supplementary Table 2**, respectively. The fact that object representations emerged at similar initial latencies for both challenge and control images (i.e. 90 and 80 ms) supports our prediction. However, the rise in decoding accuracy was slower for challenge compared to control images, with a significant decoding difference emerging at 90 ms. Despite this early lag, both curves peaked at comparable latencies, and no significant differences were observed from 220ms onwards. This temporal pattern suggests that object representations emerge more slowly for challenge than for control images, consistent with the engagement of additional recurrent processing before converging onto similar dynamics.

**Figure 4.**
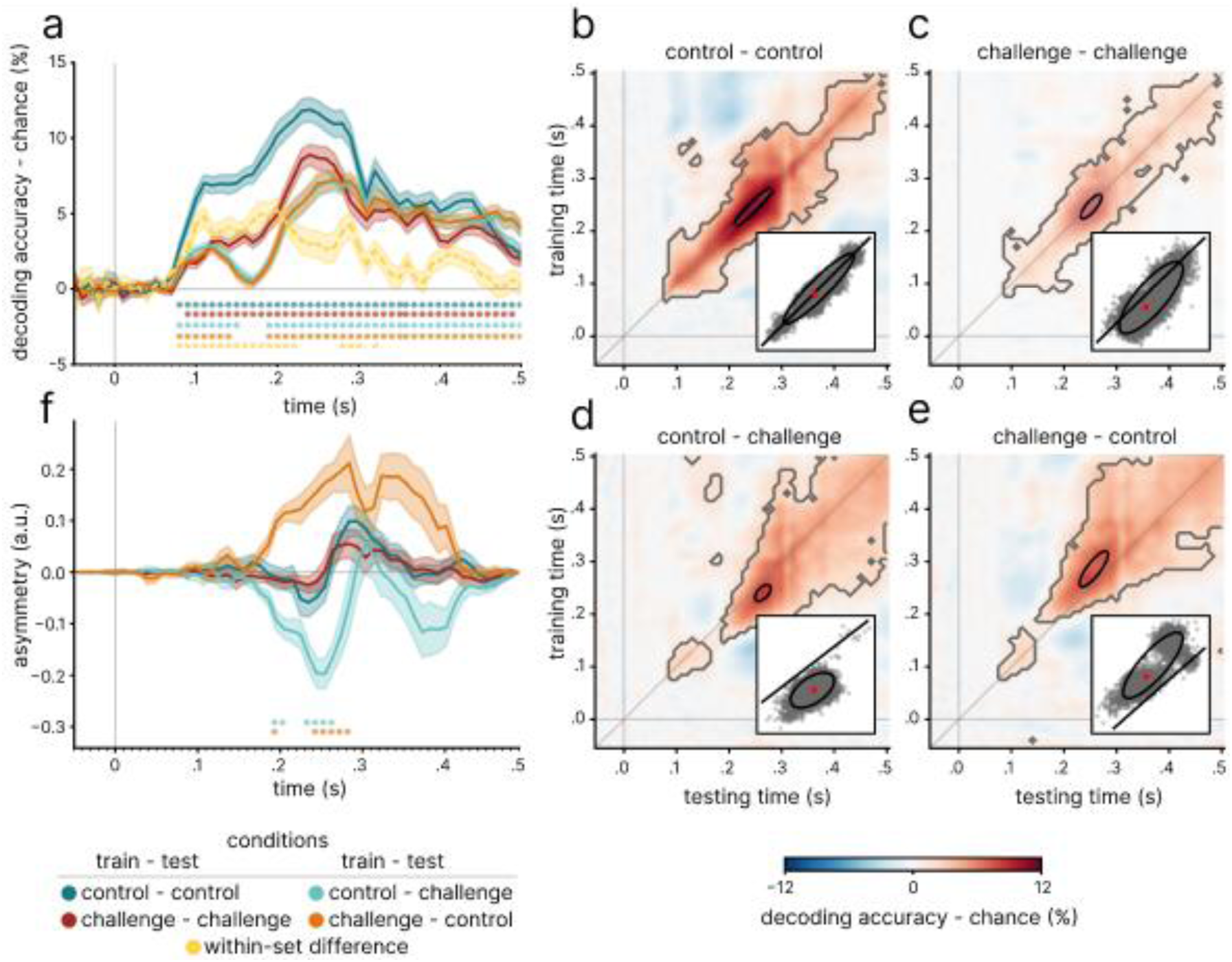
Temporal dynamics of object recognition according to decoding analysis. **a.** Time courses of decoding accuracy for control (color-coded blue) and challenge (red) images, as well as for cross-decoding from control to challenge (cyan) and from challenge to control (orange). Thick lines show subject-averaged decoding accuracy; shaded regions indicate bootstrapped SEM. The yellow line depicts the within-set decoding difference (control minus challenge). Dots below the x-axis are color-coded as time courses, and mark significant time points (sign-flip permutation test across subjects, Bonferroni-corrected across time; *p* < 0.05). **b–e.** Temporal generalization matrices (TGMs), indicates decoding accuracy for object decoding across training→testing conditions: b) control→control, c) challenge→challenge, d) control→challenge, e) challenge→control. Grey outlines denote clusters of significant decoding (sign-flip permutation tests, Bonferroni-corrected, p < 0.05). Peak decoding is marked by black 95% confidence contours. Insets show the distribution of individual peak locations (grey points), the confidence ellipse, and the group-level mean (red point). **f.** Temporally resolved antidiagonal asymmetry for each TGM (b–e), quantifying differences in decoding strength above vs. below the diagonal for each time point. Positive values reflect stronger upper-segment decoding (later training, earlier testing). Color coding follows panel a. Significant time points are marked below the x-axis (sign-flip permutation test, Bonferroni-corrected, p < 0.05). a.u. = arbitrary units.

To directly assess whether underlying object representations differ over time, we performed cross-decoding of object identity across challenge and control images (**Fig. 4a**, cyan and orange traces for each training direction; see **Supplementary Table 3** and **Supplementary Table 4** for peaks magnitudes and latencies respectively, along with 95% confidence intervals). Cross-decoding was significant from 90ms and rose sharply thereafter, closely tracking the within-condition decoding curves, and reaching a first peak at 116ms. However, between 140–190ms, this rise was disrupted by a period of non-significant cross-decoding, followed by a second peak at 273 ms, thereafter converging again with the within-condition decoding profile. This transient breakdown in cross-decoding suggests that object representations diverge across conditions at an intermediate processing phase. This could reflect the emergence of distinct object representations for challenge and control images, a temporal misalignment of otherwise similar representations, or a combination of both factors unfolding over time.

To test these possibilities, we applied a cross-condition variant of temporal generalization analysis (TGA ^43^) to characterize the temporal relationship between object representations evoked by challenge and control images. Specifically, we decoded object identity across all pairwise time points within the epoch, capturing how representations generalize over time across conditions. In this analysis, classifiers were trained on challenge images and tested on control, and vice versa. We assessed statistical significance using sign-permutation tests (1,000,000 permutations, *p* < 0.05, Bonferroni-corrected across time points). Cross-condition TGA revealed commonalities and differences in the emergence of object representations over time, delineating three stages of processing (**Fig. 4d,e; Supplementary Table 5** for peak latencies). First, an early phase centered on the diagonal (< 90 ms) indicated comparable dynamics with which similar object representations emerge for challenge and control images. Given its rapid timing, this phase likely reflects feed-forward processing ^9,44^. Second, an intermediate stage (∼140–190 ms) showed a disruption of cross-condition generalization, despite robust object information for both conditions (see **Fig. 4b,c**), indicating divergent representational trajectories. Third, a later stage (∼190–500 ms) showed asymmetric generalization adjacent to the diagonal, consistent with a delayed emergence of similar representations for challenge images. We verified the statistical significance of this delay in two complimentary ways. First, we computed an asymmetry index quantifying the differences in decoding accuracy between the upper and lower halves of antidiagonal vectors relative to each diagonal time point (**Fig. 4f**). This analysis revealed a significant asymmetry for cross-decoding TGMs between approximately 190 and 280 ms post-stimulus (100,000 sign-flip permutations across subjects, *p* < .05, Bonferroni-corrected across time points). Second, we determined the position of peak decoding in the TGMs. The peaks in the cross-decoding TGMs were located off-diagonal: peaks were found at the time point combinations [241ms, 265 ms] and [274 ms, 251 ms] respectively. In both cases, the differences of 24 ms and 23 ms were significant (*p* < .05, sign-permutation tests, 10,000 permutations; insets in **Fig. 4d,e**; **Table 5**). For completeness, we also performed TGA for both challenge and control images separately (**Fig. 4b,c**). As expected, both conditions showed symmetric generalization patterns centered on the diagonal. The asymmetry index (**Fig. 4f**) was low and not significant, and the peaks for challenge and control TGMs were on or near diagonal (challenge; [240 ms, 245 ms] control: [239 ms, 242 ms], insets in **Fig. 4b,c**; **Table 5**); their differences of 5ms and 3ms were not significant.

Together, these results indicate that neural responses to challenge and control images initially follow similar feedforward dynamics and then diverge before reconverging onto similar representations. For challenge images, this convergence is delayed, consistent with the additional time required for recurrent processing.

## Discussion

We hypothesized that when object recognition is challenging, feedforward processing within the ventral visual stream is insufficient, and additional neural resources beyond this pathway need to be recruited to implement recurrent computations. Our results support this hypothesis, showing rapid recruitment of a distributed cortical network encoding object information for images recognized accurately by humans but misclassified by a canonical feedforward model lacking recurrence. Our fMRI results revealed that these additional resources span regions across frontal and parietal regions, and are accompanied by a decorrelation of object representations across the ventral stream, which is consistent with a functional reconfiguration of the network. Our EEG data further showed that object representations for challenge images emerged with a delay, preceded by a transient processing phase (∼140-190 ms) reflecting rapid, condition-specific engagement of additional cortical resources. Together, these findings support the view that when processing demands exceed the capacity of the ventral visual stream, object recognition relies on fast recurrent computation supported by adaptively recruited frontoparietal systems.

### Cortex-wide object representations beyond the ventral stream for challenge images indicates automatic and adaptive engagement of additional resources

We found that fMRI decoding revealed object representations in frontal and parietal regions beyond the ventral visual stream, but only for challenge images. In contrast, control images elicited object representations confined to the ventral visual stream. These results reconcile two seemingly opposing accounts of object recognition: one restricting it to ventral feedforward processing ^1,2,15,28,45^, the other emphasizing broader cortical networks. This suggests that the neural machinery supporting object recognition is dynamic and depends on the sufficiency of feedforward processing within the ventral visual stream. Of note, all regions identified beyond the visual stream are anatomically connected to visual cortex ^46–50^ and contain visually responsive neurons ^18–20,51–54^. Importantly, our results go beyond establishing activity for challenge images: they show that these regions explicitly encode object identity. This indicates a direct computational role rather than a purely modulatory influence driven by factors as attention or task demands ^28,55–57^.

What is the exact role for each part of the identified cortical machinery engaged for challenge images? The network additionally engaged for challenge images spans regions frequently implicated in attention, working memory, and cognitive control, overlapping with canonical large-scale networks. Several parietal, insular, and frontal regions (iPC, IFJ, AI/OpC, PMC, v/dlPFC) are implicated in top-down attentional control, overlapping with components of the dorsal attention ^58,59^ and salience networks ^60,61^. These systems coordinate the selection of behaviorally relevant stimuli and bias sensory processing accordingly, including object-based attention in visual regions ^62–65^. In working memory, midline, temporal and frontal regions (mCC, PCC/PCu, mTC, AI/OpC, mPFC, vlPFC), overlapping with the cingulo-opercular ^66^ and default mode ^67^ networks, have been linked to the maintenance ^68,69^, prioritization, and updating of task-relevant representations ^42,70,71^. These regions are thought to support the buffering and manipulation of perceptual information when bottom-up signals are insufficient ^72–75^. Interactions with IT have also been shown during retrieval supporting associative memory ^76–78^. In cognitive control, prefrontal and parietal regions (dlPFC, vlPFC, mPFC, IFJ, AI/OpC, iPC) are linked to the frontoparietal control network ^60,79,80^, contributing to faculties that involve flexible selection, coordination, and integration of information in a goal-dependent manner ^68,69,81–83^. These different systems dynamically interact ^84–86^, and their joint recruitment here may reflect distributed, adaptive coordination in response to perceptual demands.

While we acknowledge that our findings are correlative, and cannot fully establish a causal involvement of the identified regions in object recognition, they provide a set of clearly defined targets for future causal interventions, such as using TMS in humans. In line with these claims, nonetheless, Kar & DiCarlo ^17^ have demonstrated a causal role in shaping IT responses for at least one of the regions investigated here (vlPFC), using an equivalent research approach in non-human primates.

### Visual cortical networks reorganize their representational structure for challenging object processing

Our results provide three key insights into the configuration of cortical networks during object processing. First, under challenging conditions (when additional resources are engaged) the visual network exhibits a reconfiguration, expressed in a reduced similarity between the representational geometries of higher and lower visual regions. A candidate mechanism to explain this phenomenon is increased iterative recurrent processing within the visual hierarchy, which would be required to solve object identity in challenge images, thus resulting in a representational divergence. This interpretation aligns with the observation that adding layers in a ffCNN, simulating temporal unrolling in a recurrent CNN ^87,88^, reduces similarity across layers ^89,90^; and with evidence for ubiquitous feedback effects from higher to lower visual regions during visual processing ^91–100^. An alternative mechanism is that extra-visual regions influence visual processing. For example, circuits in frontal cortex can dynamically adjust connectivity to form flexible associative mappings with feature-selective neurons ^18,42^. Notably, the reconfiguration of visual cortex was accompanied by network-wide distribution of object representations across frontal and parietal regions. Our results provide new empirical access points to test these hypotheses, for example through chronometric brain stimulation ^101,102^.

Second, irrespective of any additional resources engaged, similar object representations emerged only in high level ventral visual cortex. This pattern is consistent with higher-level ventral areas serving as a stable locus of explicit, invariant object representations ^10,32,34^ and a plausible read out source for downstream cognitive processes. However, this finding does not pinpoint the readout locus nor it falsifies the hypothesis of flexible or distributed readout sites across visual cortex, depending on behavioral relevance and task demands ^100,103–105^.

Finally, for challenge images, representational geometries in dlPFC and higher-level visual regions are negatively correlated. Given the role of dlPFC in integrating sensory evidence to guide decisions ^106–108^, the negative correlation in representational geometry suggests an antagonistic, or corrective role for dlPFC on high-level visual cortex in resolving mismatched object information, facilitating disambiguation of unclear configurations ^41,42^ or refining categorical boundaries ^19,20^. A testable prediction of this idea is that object representations in higher-level visual cortex evolve over time, from an initial state aligned with dlPFC to one anticorrelated, as recurrent processing reshapes the representation. Direct test of this hypothesis will require invasive recordings.

### Dynamics of object representations suggest rapid and transient engagement of additional cortical resources

The temporal dynamics with which object representations emerge for challenge versus control images revealed a nuanced three-partite view. In the early stage (90–140 ms), object representations were similar across image sets, indicating primarily shared initial feed-forward processing. The intermediate stage (140–190 ms) was instead characterized by lack of evidence for similar representations, despite the presence of robust information for both challenge and control images. This indicates divergent neural processing and the rapid, transient engagement of additional resources. In late stage (post 190 ms), representations realigned for challenge and control images, albeit initially delayed for challenge images. This delay mirrors previous findings showing delayed emergence of object representations when object recognition is challenging, as under occlusion, or clutter ^16,21–24,56^. The most parsimonious explanation for this delay is that it reflects additional processing time required by the intermediate stage. Taking our fMRI results into account, this results pattern suggests that the additional cortical resources engaged by challenge images are the loci of activity during this intermediate stage, before converging on common representations in higher-level visual regions.

### Limitations

Several limitations of the present study should be noted. First, our findings are correlational: while fMRI and EEG reveal spatial and temporal patterns consistent with adaptive recurrence, they cannot categorically establish a causal chain. Although prior nonhuman primate work has shown a causal role for prefrontal recurrence in shaping ventral stream responses ^17^, direct perturbation methods (e.g., TMS, pharmacological interventions, or intracranial recordings) will be needed to test the necessity of the identified regions in humans. Second, our operationalization of *challenge* and *control* images relied on the performance of a canonical feedforward CNN (AlexNet). While this provides a principled proxy for ventral feedforward processing, more recent architectures vary in their alignment with human recognition and may shift the boundary between solvable and challenging conditions. Nevertheless, the approach offers a systematic way to dissociate conditions where feedforward suffices from those requiring additional resources. Finally, although we observed recruitment of a broad frontoparietal network, our study does not resolve the division of labor across its constituent regions. Distinguishing between attentional control, working memory, and decision-related contributions will require targeted manipulations and connectivity analyses.

## Conclusion

Visual object recognition is a core cognitive function, yet the scope of its underlying neural machinery remains under debate ^45,109^. Taken together, our findings support the adaptive recurrence hypothesis, proposing that the neural machinery is flexibly configured on demand: when feedforward processing in the ventral visual stream is insufficient, additional resources across dedicated cortical networks are recruited. This engagement is rapid and transient, changing the network configuration of the ventral visual stream, and leading to the delayed emergence of object representations in higher-level ventral visual regions.

## Methods

### Participants

We conducted two separate experiments with no overlapping participants: an fMRI and an EEG experiment. All participants had normal or corrected-to-normal vision. Thirty-one participants took part in the fMRI study (N=31, mean age 27 ± 4.8 years, 21 female). Thirty-five participants took part in the EEG study (N=35, mean age 23.5 ± 4.1 years, 28 female). All participants provided their written informed consent before taking part in the study and were compensated with a either monetary sum or course credits. The experiments were approved by the local ethics committee of the Department of Education and Psychology of the Freie Universität Berlin and were conducted in accordance with the Declaration of Helsinki. In addition, we used data from a behavioral experiment comprising 88 participants on Amazon Mechanical Turk, previously published in Kar et al. ^24^.

### Experimental Stimuli

We used a set of 1,320 images from Kar et al.^24^, each depicting one of ten possible objects. These images included both photographs (sourced from MS COCO ^25^) and synthetic objects rendered with a combination of transformations (i.e., size, rotation, and position) against a natural background. This set was chosen because it spans a broad object space with varied depictions, covering a range of recognition difficulty, including images misclassified by feedforward models.

### Human Behavior dataset

To obtain human recognition performance for each image, we used behavioral data from Kar et al. ^24^, where object recognition was tested in a large-scale online experiment using Amazon Mechanical Turk. In each trial, participants briefly viewed one of 1,320 images (100 ms), followed by a two-alternative forced-choice between the target object and a distractor. Each participant saw each image only once. The resulting image-level accuracy scores quantified human recognition performance, which we later compared to model outputs to define challenge and control conditions

### Behavioral performance metric

To quantify recognition performance for both humans and the feedforward model, we computed a sensitivity index (*d*′) for each image. For a given image of object *i*, the hit rate was defined as the average accuracy across binary discriminations between *i* and all non-target objects (*j* ≠ *i*). The false alarm rate reflected the mean rate at which *i* was incorrectly chosen as the target when absent. These rates were converted into a per-image sensitivity score

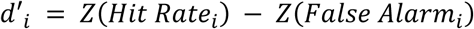

where *Z* is the inverse cumulative Gaussian. This metric was applied identically to human and model behavior, enabling a direct comparison of per-image recognition performance. For full details, see Kar et al. ^24^.

### Selection of challenge and control images

We contrasted human and model (i.e. Alexnet ^26^) *d*′ scores across the full stimulus set (1,320 images) to classify images according to feedforward solvability. Challenge images were defined as those for which human performance exceeded Alexnet’s by more than 1.5 *d*′, while control images were those with an absolute difference not exceeding 0.4 *d*′.

We then created a final, smaller stimulus selection for EEG and fMRI experiments by pairing each challenge image with the control image best matched for human *d*′. We used the Kuhn-Munkres algorithm to solve this as a linear assignment problem: finding the optimal one-to-one pairings that minimized the total difference in human *d*′ between the two sets. This ensured that any observed effects in brain signals could not be attributed to differences in human performance between the two sets. This process yielded two sets of 121 images each, with no significant difference in human performance between them (u(120)=8155, p=0.13) but a significant difference in CNN performance (u(120)=41, p<0.001; **Fig. 1d**).

### fMRI experiment

#### fMRI experimental paradigm

We recorded brain responses to the selected challenge and control images (N = 242; 8-10 repetitions each). Each trial presented an image for 500 ms (8° visual angle), followed by a 2,500 ms blank screen. Participants were instructed to respond to a color change in the fixation cross, which occurred randomly between trials (probability 0.25). In each run, we presented a random half of the images (i.e., 121), ensuring that across every two runs, all images from both sets were presented once. This procedure resulted in a run duration of approximately 7.5 minutes. Each participant completed two sessions, each consisting of 8 to 10 experimental runs.

#### fMRI data acquisition

We collected MRI data using a 64-channel head coil on a Siemens Magnetom Prisma Fit 3T system (Siemens Medical Solutions, Erlangen, Germany). A structural image was acquired with a standard T1-weighted sequence (TR = 1.9s, TE = 2.52ms, number of slices = 176, FOV = 256mm, voxel size = 1.0mm isotropic, flip angle = 9°). Functional images for the main experiment were acquired using a gradient-echo echo-planar imaging (GRE-EPI) T2*-weighted sequence with multiband acceleration factor 3, providing whole-brain coverage (TR = 1 s, TE = 33.3 ms, number of slices = 39, voxel size = 2.49 x 2.49 mm, matrix size = 82×82, FOV = 204 mm, flip angle = 70°, slice thickness = 2.5 mm, interleaved acquisition order, inter-slice gap = 0.25mm).

### fMRI data preprocessing

We preprocessed the fMRI data using SPM12 (*SPM12 Software – Statistical Parametric Mapping*, n.d.) and custom scripts in MATLAB R2021a (www.mathworks.com). First, we realigned all functional images to the first image of each run, slice-time corrected, and co-registered them to the anatomical image. Next, we estimated noise components using the aCompCor method ^110^, as implemented in the TAPAS PhysIO toolbox ^111^. This approach utilized tissue probability maps of white matter and cerebrospinal fluid derived from each participant’s anatomical image.

### Estimation of image-specific responses

To extract single-image activation patterns for decoding (**Fig. 2**), we modeled the BOLD response to each image using a general linear model (GLM), estimated separately for each run. Each image onsets and duration were included as regressors in the model and convolved with a hemodynamic response function (HRF). Since each run consisted in the presentation of a random half of the images, the model resulted in 121 regressors per run. As nuisance regressors, we included the noise components computed from the white matter and cerebrospinal fluid, as well as movement parameters with their first- and second-order derivatives.

To account for nonlinear properties in the hemodynamic response in the context of fast fMRI ^112^, we employed an HRF-fitting procedure following Prince et al. ^113^. The GLM fitting was repeated 20 times, with all regressors of interest convolved with different HRFs from an open-source library derived from the Natural Scenes Dataset ^114^. For each voxel, beta estimates for the image regressors were derived from the HRF that minimized the mean residual for that voxel. Importantly, this procedure is focused on optimizing the overall fit to the data without relying on condition-specific information, preventing the introduction of positive bias into multivariate decoding analyses. For the searchlight analyses we normalized these maps to the MNI template brain space.

### Definition of Regions of Interest (ROIs)

To perform a spatial analysis focused on different ROIs. We defined 11 key areas based on the Human Connectome Project (HCP) atlas ^27^, including visual and frontal regions. Specifically, we focused on the early visual cortex (EVC; V1-3), V4, dorsal visual areas (POC; V3a, V3b, V6, V6a, and intraparietal sulcus), inferotemporal cortex (IT; V8, fusiform face complex, posterior inferotemporal complex, ventral visual complex, and ventromedial visual areas 1-3), lateral occipital cortex (LOC; V3CD, V4t, lateral occipital areas 1-3, middle temporal area, medial superior temporal area, fundus of the superior temporal sulcus, and PH), premotor cortex (PMC; frontal eye fields, premotor eye field, Brodmann areas 6 and 55b), medial temporal cortex (mTC; parahippocampal areas 1-3 and TF, perirhinal cortex, entorhinal cortex, presubiculum, and hippocampus), medial prefrontal cortex (mPFC; posterior OFC complex, Brodmann areas 8b medial, 9 medial, 10 ventral and rostral, 24 anterior and posterior, 25, 32, and 33), orbitofrontal cortex (OFC; orbitofrontal complex, Brodmann areas 10 dorsal and posterior, 11, 13, 47 medial and superior), ventrolateral prefrontal cortex (vlPFC; inferior frontal junction, inferior frontal sulcus, Brodmann areas 44, 45, 47 lateral and rostral), and dorsolateral prefrontal cortex (dlPFC; superior frontal language area, Brodmann areas 8a, 8b lateral, 8c, 9 anterior and posterior, 46, and transitional areas 6-8). Next, we selected the 500 most activated voxels in each ROI for each participant based on t-values from the contrast stimuli > baseline, resulting in participant-specific ROI definitions.

### EEG experiment

#### EEG experimental paradigm

We presented the selected challenge and control images (N = 242; 68 repetitions each) using a rapid serial visual presentation paradigm ^115^. Sequences of 14 images (8° of visual angle) were presented in random order (200 ms on, 100 ms off), with some sequences including a distractor image (a paper clip, probability 0.56). Participants reported the presence of the distractor after the train concluded. Each participant completed two sessions of 14 runs, with each run consisting of 50 image sequences, resulting in a total session duration of approximately one hour.

#### EEG data acquisition

EEG responses to each image were recorded using a 64-channel EASYCAP system and a Brainvision actiCHamp amplifier, with a sampling rate of 1,000 Hz (Brain Products, Gilching, Germany). The electrodes were arranged according to the 10-10 system and referenced to Fz.

#### EEG data preprocessing

After acquisition, we re-referenced the signal to the mastoids to enhance the signal quality in the frontal electrodes. We then downsampled the data to 200 Hz and applied bandpass filtering between 0.1 and 40 Hz. Next, we epoched the trials from -50 ms to +500 ms around stimulus onset and performed baseline correction by subtracting the average signal in the pre-stimulus period from each epoch and channel independently. Finally, we applied multivariate noise normalization to each participant’s dataset separately ^116^. No additional artifact correction procedures were used, following standard practice in multivariate EEG decoding ^117–119^.

### Spatially resolved multivariate fMRI analysis

#### ROI-based classification of objects from fMRI data

To determine spatial differences in object representations between challenge and control images, we performed multivariate decoding analyses ^120^, using beta coefficients estimated from the GLM (see “Estimation of image-specific responses”) as input features. For each ROI, we first averaged beta estimates across repetitions of each image, yielding a single activation vector per image. To improve reliability, we constructed pseudo-trials by randomly grouping activation vectors for each object into three subsets and averaging the vectors within each subset. This procedure reduces noise while preserving object-level information and was applied separately for the control and challenge image sets. We then trained (linear) support vector machine (SVM) classifiers ^121^ on two of the three pseudo-trials per object and tested on the held-out pair, ensuring balanced train-test splits. In this classification problem, each data instance (i.e., each row of the input matrix) corresponded to a pseudo-trial: a vector of voxel activation values for a given object within an ROI. Classification was performed separately for the challenge and control sets. To estimate both within-set and cross-set decoding accuracy, classifiers trained on one set were tested on both the same and the alternate set. This procedure was repeated 100 times per object pair, and accuracies were averaged across all pairwise comparisons to yield a general index of object information for each ROI.

#### Searchlight classification of objects from fMRI data

To further examine the spatial distribution of object information comprehensively across the brain, we conducted a searchlight decoding analysis ^36,37^, using the whole-brain normalized activation maps in MNI space ^38^. For this, we defined a sphere with a 4-voxel radius around each voxel and extracted all beta values within this sphere to create a feature vector for classification. We applied the same decoding scheme used for the ROIs to each feature vector. This process was repeated for each valid voxel across the brain, resulting in classification accuracies for every voxel. Finally, these results were smoothed (FWHM=5) before statistical assessment.

#### Representational similarity analysis (RSA) across ROIs

To evaluate the similarity of object representations across different brain areas, we conducted a representational similarity analysis (RSA ^40^) using fMRI data from all previously defined ROIs. This approach allowed us to characterize the representational geometry of objects in each brain region and assess the congruence between them. For each ROI, we constructed an object-by-object representational dissimilarity matrix (RDM) based on decoding output weighted by the classifier decision values (*DV*) for each prediction. This approach implicitly emphasizes informative features, yielding a more sensitive estimate of representational dissimilarity.

Raw decoding accuracy is a limited metric, as it discretizes decisions and discards continuous dissimilarity information ^122^. To retain this information, we computed dissimilarity scores by weighting each prediction with the corresponding *DV*, reflecting the classifier’s confidence ^116^. Specifically, we used the following dissimilarity definition:

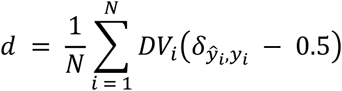

Where *N* is number of predictions, δ_*ŷi*,*yi*_ = 1 if the predicted label *ŷ*_*i*_ matches the true label *y*_*i*_, and 0 otherwise. Subtracting 0.5 centers the estimate at chance level.

In summary, computing an RDM is based on running the standard decoding analysis scheme, but, instead of collapsing all the errors into a single metric of accuracy, we retain the actual decision values between each pair of stimuli. To measure the similarity between ROIs, we then correlated the RDMs from different ROIs using the Spearman correlation coefficient.

#### Temporally resolved classification of objects from EEG data

To compare the temporal dynamics of object information in and across both sets of stimuli, we performed time-resolved decoding on the EEG data. We applied pairwise classification of object identity using linear SVM classifiers. Decoding was conducted separately within each condition (i.e., control and challenge) and across conditions (cross-decoding).

To construct robust input features, we first averaged EEG signals across repetitions of each image, yielding one time series per image. For each object pair, we randomly selected an equal number of image-specific time series per class and grouped them into three equal-sized subsets. Each subset was then averaged across trials to form a pseudo-trial, improving signal-to-noise ratio. This procedure was applied identically to both image sets. At each time point, we applied a stratified cross-validation scheme, training on two of three pseudo-trials per object and testing on the held-out pair, to maintain class balance. We repeated this process 100 times per object pair with different random assignments. Decoding accuracies were then averaged across all object pairs and repetitions to yield a temporal profile of object information. Cross-decoding was performed identically, but training and testing were conducted on different image sets.

### Temporal Generalization of EEG-Derived Object Classifiers

To assess the temporal dynamics and stability of object-related neural patterns, we performed a temporal generalization analysis on the EEG data ^43^. At each time point, we trained linear SVM classifiers and tested them across all time points in the trial, producing a temporal generalization matrix (TGM) with training times on one axis and testing times on the other. As before, this analysis was conducted both within and between image sets to evaluate the temporal specificity and generalizability of object representations over time and stimulus sets.

### Symmetry quantification of temporal generalization matrices

We computed a temporally resolved asymmetry index to compare decoding accuracies above and below the diagonal of the TGM. For each time point *t*, we extracted the antidiagonal centered at (*t*, *t*); that is, the set of entries orthogonal to the main diagonal and passing through (*t*, *t*). Each antidiagonal was split into two parts: an upper segment *U*_*t*_ containing entries above the diagonal, and a lower segment *L*_*t*_ containing the entries below. To improve signal-to-noise ratio, only statistically significant entries were included. The asymmetry index at time *t*, denoted *A*_*t*_, was computed as

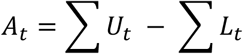

This yielded a time series of asymmetry scores *A* = {*A*_*t*_}, quantifying the directional imbalance in generalization patterns relative to the diagonal at each time point. We applied a bootstrap procedure with 10,000 resampling iterations across subjects and computed *A* for each repetition. In every iteration, subjects were resampled with replacement, preserving the original sample size.

#### Statistical hypothesis testing

To assess the statistical significance of our results, we employed non-parametric one-sample permutation tests with random sign flips, permuted across subjects. The sign-flipping procedure involved randomly inverting the sign of subject-level data. This process was repeated 10,000 times to produce the null distributions. These distributions were then used to estimate the probability of observing the empirical value under the null hypothesis. For decoding analysis, the test was one-sided, comparing metrics to chance level, while for RSA, it was two-sided, comparing metrics to zero. All results were corrected for multiple comparisons within each analysis using the Bonferroni method, with exception of searchlight decoding analysis on fMRI data. In that case, we used Benjamini-Yekutieli procedure ^39^, which controls the false discovery rate under arbitrary dependence. This is a necessary consideration for spatially-correlated voxel data.

## Acknowledgements

PO was supported by the Scholarship Program of the National Agency for Research and Development (2019-72200281). R.M.C. is supported by German Research Council (DFG) grants (CI 241/1-1, CI 241/1-3, CI 241/1-7 and INST 272/297-1), the European Research Council (ERC) starting grant (ERC-StG-2018-803370) and the ERC Consolidator grant (ERC-CoG-2024101123101). DV is supported by a Novo Nordisk Foundation Emerging Investigator Fellowship (NNF19OC-0054895) and an ERC Starting Grant (ERC-StG-2019–850404).

## Supplementary information

**Supplementary Table 1.** Data underlying Figure 4a: Peak decoding accuracy for within-set decoding.

| Condition | Peak accuracy (%) |  |  |
| --- | --- | --- | --- |
|  | Mean | CI lower | CI upper |
| Control – control | 12.03 | 10.51 | 13.43 |
| Challenge - challenge | 9.03 | 7.68 | 10.35 |

Mean peak decoding accuracy above chance (%), with 95% bootstrap confidence intervals (CI), for control-control and challenge-challenge conditions

**Supplementary Table 2.** Data underlying Figure 4a: Peak decoding latency for within-set decoding.

| Condition | Peak latency (seconds) |  |  |
| --- | --- | --- | --- |
|  | Mean | CI lower | CI upper |
| Control – control | .241 | .210 | .280 |
| Challenge – challenge | .246 | .230 | .270 |

Mean latency of peak decoding (seconds), with 95% bootstrap confidence intervals (CI), for control-control and challenge-challenge conditions

**Supplementary Table 3.** Data underlying Figure 4a: Peak decoding accuracy for cross-set decoding.

| Condition | Peak accuracy (%) |  |  |  |  |  |
| --- | --- | --- | --- | --- | --- | --- |
|  | Peak A |  |  | Peak B |  |  |
|  | Mean | CI lower | CI upper | Mean | CI lower | CI upper |
| Control – challenge | 2.92 | 2.43 | 3.47 | 7.28 | 6.16 | 8.47 |
| Challenge – control | 2.68 | 2.17 | 3.23 | 7.45 | 6.33 | 8.57 |

Mean peak decoding accuracy above chance (%), with 95% bootstrap confidence intervals (CI), for control-challenge and challenge-control conditions

**Supplementary Table 4.** Data underlying Figure 4a: Peak decoding latency for cross-set decoding.

| Condition | Peak latency (seconds) |  |  |  |  |  |
| --- | --- | --- | --- | --- | --- | --- |
|  | Peak A |  |  | Peak B |  |  |
|  | Mean | CI lower | CI upper | Mean | CI lower | CI upper |
| Control – challenge | .116 | .100 | .120 | .273 | .250 | .300 |
| Challenge – control | .119 | .110 | .120 | .273 | .250 | .290 |

Mean latency of peak decoding (seconds), with 95% bootstrap confidence intervals (CI), for control-challenge and challenge-control conditions

**Supplementary Table 5.** Data underlying Figure 4b-e: Timing of temporal generalization peak accuracy.

| Condition | Timing of peak generalization accuracy (seconds) |  |  |  |  |  |
| --- | --- | --- | --- | --- | --- | --- |
|  | X axis (training time) |  |  | Y axis (testing time) |  |  |
|  | Mean | CI lower | CI upper | Mean | CI lower | CI upper |
| **Control – control** | .239 | .210 | .280 | .242 | .210 | .280 |
| **Challenge – challenge** | .240 | .230 | .270 | .248 | .240 | .270 |
| **Control – challenge** | .241 | .230 | .250 | .265 | .250 | .270 |
| **Challenge - control** | .274 | .260 | .320 | .251 | .230 | .280 |

Mean time of peak generalization accuracy (seconds), with 95% bootstrap confidence intervals (CI), for within- and across-set temporal generalization.

## References

1. DiCarlo, J. J., Zoccolan, D. & Rust, N. C. How Does the Brain Solve Visual Object Recognition? Neuron 73, 415–434 (2012).

2. Grill-Spector, K. & Weiner, K. S. The functional architecture of the ventral temporal cortex and its role in categorization. Nature Reviews Neuroscience 15, 536–548 (2014).

3. Riesenhuber, M. & Poggio, T. Hierarchical models of object recognition in cortex. Nature Neuroscience 2, 1019–1025 (1999).

4. Ungerleider, L. G. & Haxby, J. V. ‘What’ and ‘where’ in the human brain. Current Opinion in Neurobiology 4, 157–165 (1994).

5. Logothetis, N. & Sheinberg, D. Visual object recognition. Annu Rev Neurosci 19, 577–621 (1996).

6. Wallis, G. & Rolls, E. T. INVARIANT FACE AND OBJECT RECOGNITION IN THE VISUAL SYSTEM. Progress in Neurobiology 51, 167–194 (1997).

7. Rust, N. C. & DiCarlo, J. J. Selectivity and Tolerance (“Invariance”) Both Increase as Visual Information Propagates from Cortical Area V4 to IT. J. Neurosci. 30, 12978 (2010).

8. Kar, K. & DiCarlo, J. J. The Quest for an Integrated Set of Neural Mechanisms Underlying Object Recognition in Primates. Annual Review of Vision Science 10, 91–121 (2024).

9. Thorpe, S., Fize, D. & Marlot, C. Speed of processing in the human visual system. Nature 381, 520–522 (1996).

10. Hung, C. P., Kreiman, G., Poggio, T. & DiCarlo, J. J. Fast Readout of Object Identity from Macaque Inferior Temporal Cortex. Science 310, 863–866 (2005).

11. Felleman, D. & Van Essen, D. Distributed hierarchical processing in the primate cerebral cortex. Cereb Cortex 1, 1–47 (1991).

12. Sporns, O. & Zwi, J. D. The small world of the cerebral cortex. Neuroinformatics 2, 145–162 (2004).

13. Markov, N. T. et al. Anatomy of hierarchy: Feedforward and feedback pathways in macaque visual cortex. Journal of Comparative Neurology 522, 225–259 (2014).

14. Kietzmann, T. C. et al. Recurrence is required to capture the representational dynamics of the human visual system. Proceedings of the National Academy of Sciences 116, 21854–21863 (2019).

15. Kreiman, G. & Serre, T. Beyond the feedforward sweep: feedback computations in the visual cortex. Annals of the New York Academy of Sciences 1464, 222–241 (2020).

16. Rajaei, K., Mohsenzadeh, Y., Ebrahimpour, R. & Khaligh-Razavi, S.-M. Beyond core object recognition: Recurrent processes account for object recognition under occlusion. PLOS Computational Biology 15, e1007001 (2019).

17. Kar, K. & DiCarlo, J. J. Fast Recurrent Processing via Ventrolateral Prefrontal Cortex Is Needed by the Primate Ventral Stream for Robust Core Visual Object Recognition. Neuron 109, 164–176.e5 (2021).

18. Cromer, J. A., Roy, J. E. & Miller, E. K. Representation of Multiple, Independent Categories in the Primate Prefrontal Cortex. Neuron 66, 796–807 (2010).

19. Freedman, D. J., Riesenhuber, M., Poggio, T. & Miller, E. K. Categorical Representation of Visual Stimuli in the Primate Prefrontal Cortex. Science 291, 312–316 (2001).

20. Freedman, D. J., Riesenhuber, M., Poggio, T. & Miller, E. K. A Comparison of Primate Prefrontal and Inferior Temporal Cortices during Visual Categorization. J. Neurosci. 23, 5235 (2003).

21. Graumann, M., Ciuffi, C., Dwivedi, K., Roig, G. & Cichy, R. M. The spatiotemporal neural dynamics of object location representations in the human brain. Nature Human Behaviour 6, 796–811 (2022).

22. Groen, I. I. A. et al. Scene complexity modulates degree of feedback activity during object detection in natural scenes. PLOS Computational Biology 14, e1006690 (2019).

23. Tang, H. et al. Recurrent computations for visual pattern completion. Proceedings of the National Academy of Sciences 115, 8835–8840 (2018).

24. Kar, K., Kubilius, J., Schmidt, K., Issa, E. B. & DiCarlo, J. J. Evidence that recurrent circuits are critical to the ventral stream’s execution of core object recognition behavior. Nature Neuroscience 22, 974–983 (2019).

25. Lin, T.-Y. et al. Microsoft coco: Common objects in context. in Computer vision–ECCV 2014: 13th European conference, zurich, Switzerland, September 6-12, 2014, proceedings, part v 13 740–755 (Springer, 2014).

26. Krizhevsky, A., Sutskever, I. & Hinton, G. E. Imagenet classification with deep convolutional neural networks. Advances in neural information processing systems 25, (2012).

27. Glasser, M. F. et al. A multi-modal parcellation of human cerebral cortex. Nature 536, 171– 178 (2016).

28. Kravitz, D. J., Saleem, K. S., Baker, C. I., Ungerleider, L. G. & Mishkin, M. The ventral visual pathway: an expanded neural framework for the processing of object quality. Trends in Cognitive Sciences 17, 26–49 (2013).

29. Gilbert, C. D. & Li, W. Top-down influences on visual processing. Nature Reviews Neuroscience 14, 350–363 (2013).

30. Haxby, J. V. et al. Distributed and Overlapping Representations of Faces and Objects in Ventral Temporal Cortex. Science 293, 2425–2430 (2001).

31. Norman, K. A., Polyn, S. M., Detre, G. J. & Haxby, J. V. Beyond mind-reading: multi-voxel pattern analysis of fMRI data. Trends in Cognitive Sciences 10, 424–430 (2006).

32. Grill-Spector, K., Kourtzi, Z. & Kanwisher, N. The lateral occipital complex and its role in object recognition. Vision Research 41, 1409–1422 (2001).

33. Martin, A. & Chao, L. L. Semantic memory and the brain: structure and processes. Current Opinion in Neurobiology 11, 194–201 (2001).

34. Majaj, N. J., Hong, H., Solomon, E. A. & DiCarlo, J. J. Simple Learned Weighted Sums of Inferior Temporal Neuronal Firing Rates Accurately Predict Human Core Object Recognition Performance. J. Neurosci. 35, 13402 (2015).

35. Rust, N. C. & DiCarlo, J. J. Balanced Increases in Selectivity and Tolerance Produce Constant Sparseness along the Ventral Visual Stream. J. Neurosci. 32, 10170 (2012).

36. Kriegeskorte, N., Goebel, R. & Bandettini, P. Information-based functional brain mapping. Proceedings of the National Academy of Sciences 103, 3863–3868 (2006).

37. Allefeld, C. & Haynes, J.-D. Searchlight-based multi-voxel pattern analysis of fMRI by cross-validated MANOVA. NeuroImage 89, 345–357 (2014).

38. Mazziotta, J. et al. A Four-Dimensional Probabilistic Atlas of the Human Brain. Journal of the American Medical Informatics Association 8, 401–430 (2001).

39. Benjamini, Y. & Yekutieli, D. The Control of the False Discovery Rate in Multiple Testing under Dependency. The Annals of Statistics 29, 1165–1188 (2001).

40. Kriegeskorte, N., Mur, M. & Bandettini, P. A. Representational similarity analysis - connecting the branches of systems neuroscience. Frontiers in Systems Neuroscience Volume 2-2008, (2008).

41. Amer, T. & Davachi, L. Extra-hippocampal contributions to pattern separation. eLife 12, e82250 (2023).

42. Manohar, S. G., Zokaei, N., Fallon, S. J., Vogels, T. P. & Husain, M. Neural mechanisms of attending to items in working memory. Neuroscience & Biobehavioral Reviews 101, 1–12 (2019).

43. King, J.-R. & Dehaene, S. Characterizing the dynamics of mental representations: the temporal generalization method. Trends in Cognitive Sciences 18, 203–210 (2014).

44. Lamme, V. A. F. & Roelfsema, P. R. The distinct modes of vision offered by feedforward and recurrent processing. Trends in Neurosciences 23, 571–579 (2000).

45. van Bergen, R. S. & Kriegeskorte, N. Going in circles is the way forward: the role of recurrence in visual inference. Current Opinion in Neurobiology 65, 176–193 (2020).

46. Barbas, H. Anatomic organization of basoventral and mediodorsal visual recipient prefrontal regions in the rhesus monkey. Journal of Comparative Neurology 276, 313–342 (1988).

47. Kondo, H., Saleem, K. S. & Price, J. L. Differential connections of the temporal pole with the orbital and medial prefrontal networks in macaque monkeys. Journal of Comparative Neurology 465, 499–523 (2003).

48. Saleem, K. S., Kondo, H. & Price, J. L. Complementary circuits connecting the orbital and medial prefrontal networks with the temporal, insular, and opercular cortex in the macaque monkey. Journal of Comparative Neurology 506, 659–693 (2008).

49. Webster, M. J., Bachevalier, J. & Ungerleider, L. G. Connections of Inferior Temporal Areas TEO and TE with Parietal and Frontal Cortex in Macaque Monkeys. Cerebral Cortex 4, 470–483 (1994).

50. Yeterian, E. H., Pandya, D. N., Tomaiuolo, F. & Petrides, M. The cortical connectivity of the prefrontal cortex in the monkey brain. Cortex 48, 58–81 (2012).

51. Bar, M. A Cortical Mechanism for Triggering Top-Down Facilitation in Visual Object Recognition. Journal of Cognitive Neuroscience 15, 600–609 (2003).

52. Bar, M. et al. Top-down facilitation of visual recognition. Proceedings of the National Academy of Sciences 103, 449–454 (2006).

53. Rose, O. & Ponce, C. R. A concentration of visual cortex-like neurons in prefrontal cortex. Nature Communications 15, 7002 (2024).

54. Schendan, H. E. & Stern, C. E. Where Vision Meets Memory: Prefrontal–Posterior Networks for Visual Object Constancy during Categorization and Recognition. Cerebral Cortex 18, 1695–1711 (2008).

55. Rao, R. P. N. & Ballard, D. H. Predictive coding in the visual cortex: a functional interpretation of some extra-classical receptive-field effects. Nature Neuroscience 2, 79–87 (1999).

56. Fyall, A. M., El-Shamayleh, Y., Choi, H., Shea-Brown, E. & Pasupathy, A. Dynamic representation of partially occluded objects in primate prefrontal and visual cortex. eLife 6, e25784 (2017).

57. Konkle, T. & Alvarez, G. Cognitive steering in deep neural networks via long-range modulatory feedback connections. Advances in Neural Information Processing Systems 36, 21613– 21634 (2023).

58. Corbetta, M. & Shulman, G. L. Control of goal-directed and stimulus-driven attention in the brain. Nature Reviews Neuroscience 3, 201–215 (2002).

59. Szczepanski, S. M., Pinsk, M. A., Douglas, M. M., Kastner, S. & Saalmann, Y. B. Functional and structural architecture of the human dorsal frontoparietal attention network. Proceedings of the National Academy of Sciences 110, 15806–15811 (2013).

60. Seeley, W. W. et al. Dissociable Intrinsic Connectivity Networks for Salience Processing and Executive Control. J. Neurosci. 27, 2349 (2007).

61. Sridharan, D., Levitin, D. J. & Menon, V. A critical role for the right fronto-insular cortex in switching between central-executive and default-mode networks. Proceedings of the National Academy of Sciences 105, 12569–12574 (2008).

62. Baldauf, D. & Desimone, R. Neural Mechanisms of Object-Based Attention. Science 344, 424–427 (2014).

63. Bichot, N. P., Heard, M. T., DeGennaro, E. M. & Desimone, R. A Source for Feature-Based Attention in the Prefrontal Cortex. Neuron 88, 832–844 (2015).

64. Gregoriou, G. G., Rossi, A. F., Ungerleider, L. G. & Desimone, R. Lesions of prefrontal cortex reduce attentional modulation of neuronal responses and synchrony in V4. Nature Neuroscience 17, 1003–1011 (2014).

65. Monosov, I. E., Sheinberg, D. L. & Thompson, K. G. The Effects of Prefrontal Cortex Inactivation on Object Responses of Single Neurons in the Inferotemporal Cortex during Visual Search. J. Neurosci. 31, 15956 (2011).

66. Dosenbach, N. U. F. et al. A Core System for the Implementation of Task Sets. Neuron 50, 799–812 (2006).

67. Raichle, M. E. The brain’s default mode network. Annual review of neuroscience 38, 433–447 (2015).

68. Miller, E. K. & Cohen, J. D. An integrative theory of prefrontal cortex function. Annual review of neuroscience 24, 167–202 (2001).

69. Murray, J. D. et al. Stable population coding for working memory coexists with heterogeneous neural dynamics in prefrontal cortex. Proceedings of the National Academy of Sciences 114, 394–399 (2017).

70. Menon, V. & Uddin, L. Q. Saliency, switching, attention and control: a network model of insula function. Brain Structure and Function 214, 655–667 (2010).

71. Myers, N. E., Stokes, M. G. & Nobre, A. C. Prioritizing Information during Working Memory: Beyond Sustained Internal Attention. Trends in Cognitive Sciences 21, 449–461 (2017).

72. Panichello, M. F. & Buschman, T. J. Shared mechanisms underlie the control of working memory and attention. Nature 592, 601–605 (2021).

73. Peters, B. & Kriegeskorte, N. Capturing the objects of vision with neural networks. Nature Human Behaviour 5, 1127–1144 (2021).

74. Bays, P. M., Schneegans, S., Ma, W. J. & Brady, T. F. Representation and computation in visual working memory. Nature Human Behaviour 8, 1016–1034 (2024).

75. Van Ede, F. & Nobre, A. C. Turning attention inside out: How working memory serves behavior. Annual review of psychology 74, 137–165 (2023).

76. Kadohisa, M. et al. Frontal and temporal coding dynamics in successive steps of complex behavior. Neuron 111, 430–443.e3 (2023).

77. Miller, E. K., Erickson, C. A. & Desimone, R. Neural Mechanisms of Visual Working Memory in Prefrontal Cortex of the Macaque. J. Neurosci. 16, 5154 (1996).

78. Tomita, H., Ohbayashi, M., Nakahara, K., Hasegawa, I. & Miyashita, Y. Top-down signal from prefrontal cortex in executive control of memory retrieval. Nature 401, 699–703 (1999).

79. Duncan, J. The multiple-demand (MD) system of the primate brain: mental programs for intelligent behaviour. Trends in Cognitive Sciences 14, 172–179 (2010).

80. Dixon, M. L. et al. Heterogeneity within the frontoparietal control network and its relationship to the default and dorsal attention networks. Proceedings of the National Academy of Sciences 115, E1598–E1607 (2018).

81. Cole, M. W. et al. Multi-task connectivity reveals flexible hubs for adaptive task control. Nature Neuroscience 16, 1348–1355 (2013).

82. Stokes, M. G. et al. Dynamic Coding for Cognitive Control in Prefrontal Cortex. Neuron 78, 364–375 (2013).

83. Fedorenko, E., Duncan, J. & Kanwisher, N. Broad domain generality in focal regions of frontal and parietal cortex. Proceedings of the National Academy of Sciences 110, 16616–16621 (2013).

84. Spreng, R. N., Sepulcre, J., Turner, G. R., Stevens, W. D. & Schacter, D. L. Intrinsic Architecture Underlying the Relations among the Default, Dorsal Attention, and Frontoparietal Control Networks of the Human Brain. Journal of Cognitive Neuroscience 25, 74–86 (2013).

85. Vatansever, D., Menon, D. K. & Stamatakis, E. A. Default mode contributions to automated information processing. Proceedings of the National Academy of Sciences 114, 12821– 12826 (2017).

86. Shine, J. M. et al. The Dynamics of Functional Brain Networks: Integrated Network States during Cognitive Task Performance. Neuron 92, 544–554 (2016).

87. Miller, J. & Hardt, M. Stable recurrent models. arXiv preprint arXiv:1805.10369 (2018).

88. Nayebi, A. et al. Task-driven convolutional recurrent models of the visual system. Advances in neural information processing systems 31, (2018).

89. Kornblith, S., Norouzi, M., Lee, H. & Hinton, G. Similarity of neural network representations revisited. in International conference on machine learning 3519–3529 (PMLR, 2019).

90. Raghu, M., Unterthiner, T., Kornblith, S., Zhang, C. & Dosovitskiy, A. Do vision transformers see like convolutional neural networks? Advances in neural information processing systems 34, 12116–12128 (2021).

91. Haimerl, C., Ruff, D. A., Cohen, M. R., Savin, C. & Simoncelli, E. P. Targeted V1 comodulation supports task-adaptive sensory decisions. Nature Communications 14, 7879 (2023).

92. Hupé, J. M. et al. Cortical feedback improves discrimination between figure and background by V1, V2 and V3 neurons. Nature 394, 784–787 (1998).

93. Kornblith, S. & Tsao, D. Y. How thoughts arise from sights: inferotemporal and prefrontal contributions to vision. Current Opinion in Neurobiology 46, 208–218 (2017).

94. Lamme, V. The neurophysiology of figure-ground segregation in primary visual cortex. J. Neurosci. 15, 1605 (1995).

95. Li, W., Piëch, V. & Gilbert, C. D. Perceptual learning and top-down influences in primary visual cortex. Nature Neuroscience 7, 651–657 (2004).

96. Li, W., Piëch, V. & Gilbert, C. D. Contour Saliency in Primary Visual Cortex. Neuron 50, 951– 962 (2006).

97. Ni, A. M., Murray, S. O. & Horwitz, G. D. Object-Centered Shifts of Receptive Field Positions in Monkey Primary Visual Cortex. Current Biology 24, 1653–1658 (2014).

98. Piëch, V., Li, W., Reeke, G. N. & Gilbert, C. D. Network model of top-down influences on local gain and contextual interactions in visual cortex. Proceedings of the National Academy of Sciences 110, E4108–E4117 (2013).

99. Koivisto, M. & Silvanto, J. Visual feature binding: The critical time windows of V1/V2 and parietal activity. NeuroImage 59, 1608–1614 (2012).

100. McManus, J. N. J., Li, W. & Gilbert, C. D. Adaptive shape processing in primary visual cortex. Proceedings of the National Academy of Sciences 108, 9739–9746 (2011).

101. Pascual-Leone, A. & Walsh, V. Fast Backprojections from the Motion to the Primary Visual Area Necessary for Visual Awareness. Science 292, 510–512 (2001).

102. Sack, A. T., Camprodon, J. A., Pascual-Leone, A. & Goebel, R. The Dynamics of Interhemispheric Compensatory Processes in Mental Imagery. Science 308, 702–704 (2005).

103. Contier, O., Baker, C. I. & Hebart, M. N. Distributed representations of behaviour-derived object dimensions in the human visual system. Nature Human Behaviour 8, 2179–2193 (2024).

104. Jagadeesh, A. V. & Gardner, J. L. V1-and IT-like representations are directly accessible to human visual perception. in SVRHM 2021 Workshop@ NeurIPS (2021).

105. Kang, I. & Maunsell, J. H. R. The Correlation of Neuronal Signals with Behavior at Different Levels of Visual Cortex and Their Relative Reliability for Behavioral Decisions. J. Neurosci. 40, 3751 (2020).

106. Hanks, T. D. & Summerfield, C. Perceptual Decision Making in Rodents, Monkeys, and Humans. Neuron 93, 15–31 (2017).

107. Philiastides, M. G., Auksztulewicz, R., Heekeren, H. R. & Blankenburg, F. Causal Role of Dorsolateral Prefrontal Cortex in Human Perceptual Decision Making. Current Biology 21, 980–983 (2011).

108. Heekeren, H. R., Marrett, S., Bandettini, P. A. & Ungerleider, L. G. A general mechanism for perceptual decision-making in the human brain. Nature 431, 859–862 (2004).

109. Peters, B. et al. How does the primate brain combine generative and discriminative computations in vision? ArXiv arXiv-2401 (2024).

110. Behzadi, Y., Restom, K., Liau, J. & Liu, T. T. A component based noise correction method (CompCor) for BOLD and perfusion based fMRI. NeuroImage 37, 90–101 (2007).

111. Kasper, L. et al. The PhysIO Toolbox for Modeling Physiological Noise in fMRI Data. Journal of Neuroscience Methods 276, 56–72 (2017).

112. Polimeni, J. R. & Lewis, L. D. Imaging faster neural dynamics with fast fMRI: A need for updated models of the hemodynamic response. Progress in Neurobiology 207, 102174 (2021).

113. Prince, J. S. et al. Improving the accuracy of single-trial fMRI response estimates using GLMsingle. eLife 11, e77599 (2022).

114. Allen, E. J. et al. A massive 7T fMRI dataset to bridge cognitive neuroscience and artificial intelligence. Nature Neuroscience 25, 116–126 (2022).

115. Grootswagers, T., Robinson, A. K. & Carlson, T. A. The representational dynamics of visual objects in rapid serial visual processing streams. NeuroImage 188, 668–679 (2019).

116. Guggenmos, M., Sterzer, P. & Cichy, R. M. Multivariate pattern analysis for MEG: A comparison of dissimilarity measures. NeuroImage 173, 434–447 (2018).

117. Grootswagers, T., Wardle, S. G. & Carlson, T. A. Decoding Dynamic Brain Patterns from Evoked Responses: A Tutorial on Multivariate Pattern Analysis Applied to Time Series Neuroimaging Data. Journal of Cognitive Neuroscience 29, 677–697 (2017).

118. Delorme, A. EEG is better left alone. Scientific Reports 13, 2372 (2023).

119. Kessler, R., Enge, A. & Skeide, M. A. How EEG preprocessing shapes decoding performance. Communications Biology 8, 1039 (2025).

120. Haxby, J. V., Connolly, A. C. & Guntupalli, J. S. Decoding neural representational spaces using multivariate pattern analysis. Annual review of neuroscience 37, 435–456 (2014).

121. Cortes, C. & Vapnik, V. Support-vector networks. Machine Learning 20, 273–297 (1995).

122. Walther, A. et al. Reliability of dissimilarity measures for multi-voxel pattern analysis. NeuroImage 137, 188–200 (2016).

